# *De novo* identification of the specificities of recurrent human T cell receptors

**DOI:** 10.1101/2025.08.03.668342

**Authors:** Mithila Kasbe, Berkay Yahsi, Kalyn Whitehead, Jiwon Oh, Mashwiyat Mosharraf, Kayla Sohn, Trinh Phan, Mark N. Lee

## Abstract

T cell repertoires of different individuals occasionally converge on the same T cell receptor (TCR) sequence as a solution to target immunodominant epitopes. A complete mapping of these “public” TCR specificities may enable a global understanding of population-level immune histories. Here, we sought to determine the antigen specificities of public TCRs with unknown target identity. We developed a functional screening workflow in which we screen panels of TCRs for reactivity to individual viral genomes or to approximately 1,000 viral reference strains, and then sort out the immunogenic peptides by labeling antigen-presenting cells that are in proximity to activated T cells. Using this workflow, we identified the target specificities of T cells that are circulating in up to 14% of individuals, including a pre-COVID-19 seasonal coronavirus-reactive TCR that cross-reacts with peptides within pandemic coronaviruses; an influenza B-reactive TCR that targets a highly conserved epitope; and TCRs targeting Herpesviridae family viruses that cause long-term latent infections. Our results demonstrate an efficient strategy to reveal public T cell memories *de novo*, offering a window into shared immune exposures.

## INTRODUCTION

Memories of past viral exposures are preserved in the DNA of T cells within their recombined T cell receptor (TCR) genes, and can be read out using TCR sequencing *(1, 2)*. Decoding the target antigen(s) of each TCR sequence enables interpretation of this exposure record. A subset of TCR sequences has been recurrently found within clonally-expanded T cells in different individuals and associated with similar past exposure history, e.g. to infection by common human viruses (*1, 3*). Thus, these “public” TCR sequences are biomarkers of specific viral exposures. A complete mapping of public TCR specificities may enable a global understanding of disease exposures in each individual.

Most public TCR sequences have unknown specificities (*3*). Databases/datasets that have tracked accumulated knowledge of public TCR specificities are dominated by epitopes within cytomegalovirus (CMV), Epstein-Barr virus (EBV), influenza (Flu) A, and the pandemic coronavirus SARS-CoV-2 (*3–5)*. Beyond these viruses, public TCRs targeting many other common exposures – including seasonal (“common cold”) coronaviruses and herpes simplex viruses (HSV), etc. – are nearly absent from these databases, despite the high incidence of these infections within the human population (*6, 7*). A more complete mapping of public TCR specificities will aid in the interpretation of T cell sequencing data (*8*–*11*), as well as in the development of blood-based diagnostics (*1, 2, 12*) and anti-viral therapeutics (*13, 14*).

A major bottleneck in the effort to identify public TCR specificities has been the difficulty to test large numbers of putative viral peptides for immunogenicity due to scaling limitations of epitope identification assays that are in widespread use, including tetramers and ELISPOT (enzyme-linked immunospot) (*15, 16*). Thus, recently, high throughput T cell antigen identification assays have been developed to sort immunogenic epitopes from libraries of many thousands of encoded peptides (*17*–*20*). However, these assays are limited by the need to test single TCRs one by one. Eventually, solving the specificities of large numbers of T cells within the immune repertoire – including the set of many thousands of public TCRs (*3*) – will require approaches that preserve this highly-scaled putative antigen testing, but better scale in TCR numbers.

Here, we sought to identify the target specificities of public TCRs *de novo*. We developed a workflow, which we refer to as “AIMcap (activation-induced marker-mediated capture of target epitopes)”, in which we first screen a panel of public TCRs for reactivity to hundreds or thousands of peptides in single wells of a 96-well plate; and then, using a novel activation-induced marker (AIM)-based antigen sorting method, we identify the specific viral peptide(s) targeted within the well.

## RESULTS

### Selection of public T cell receptors using deep TCRβ and single-cell sequencing datasets

We evaluated a set of TCRβ sequences that are: (i) recurrently found in clonally-expanded T cells in the peripheral blood of healthy individuals (in samples collected pre-COVID-19 pandemic (*1*)), (ii) found in single-cell TCR sequencing datasets along with a paired TCRα gene, and (iii) recurrently found in individuals who share a common HLA class I allele (suggesting antigen recognition rather than increased generational likelihood during VDJ recombination) (*3*) (**Fig. 1, A** and **B**, and **fig. S1A**, and **tables S1** to **S3**). A subset of these T cells had evidence of specificity for CMV, EBV, or Flu A, but others have unknown target specificities (*3*) (**table S3**).

**Fig. 1.**
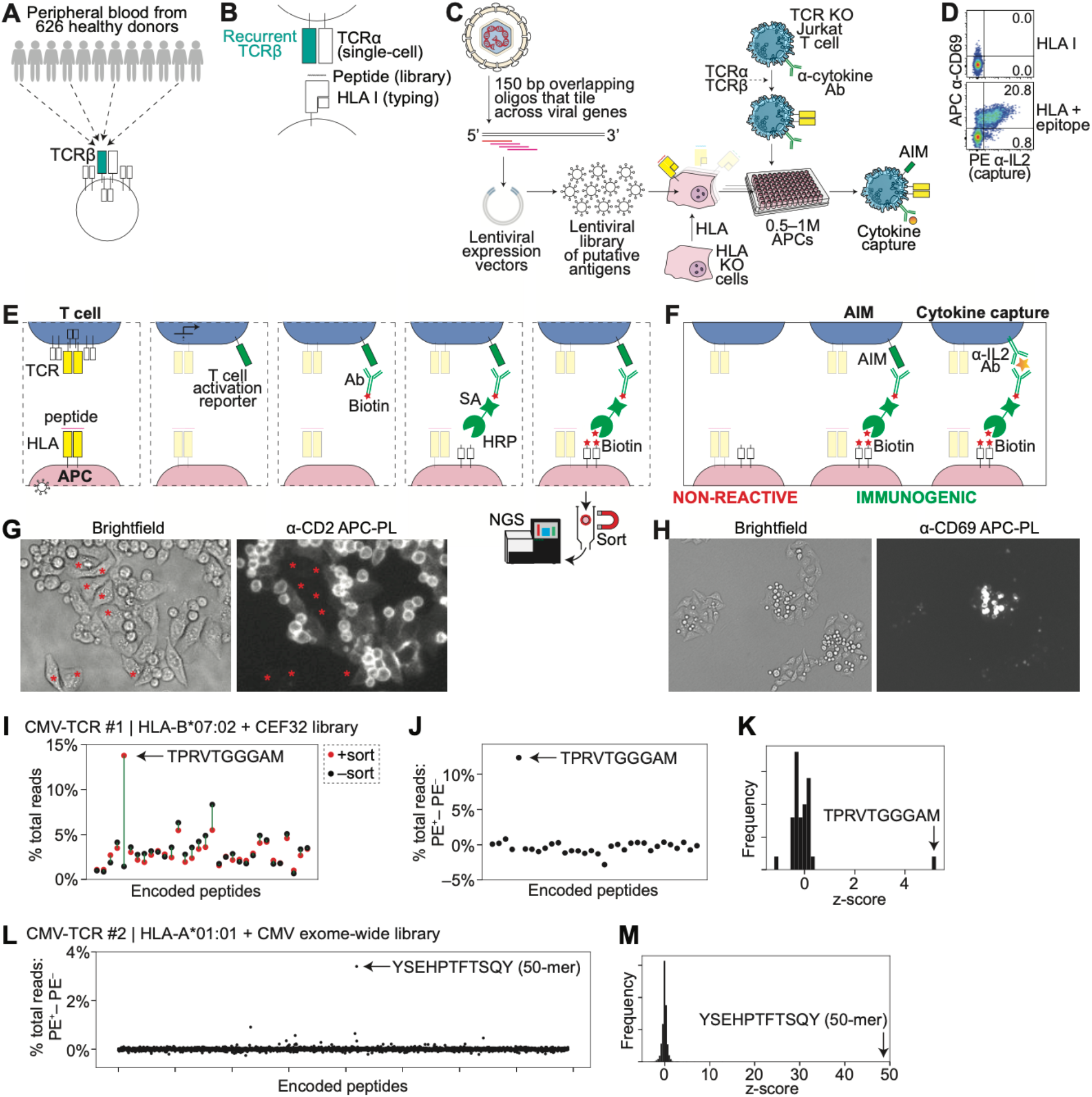
Development of the AIMcap system to identify the targets of public TCRs. (**A**) Schematic of shared (“public”) TCRβ sequences in the peripheral blood of healthy donors. (**B**) Schematic showing sources of genes used to functionally test HLA–peptide–TCR interactions. (**C**) Schematic of the system to screen viral genome libraries. Viral genes are tiled with overlapping oligos, which are cloned into lentiviral expression vectors. Lentivirus is transduced, along with specific HLA alleles, into HLA knockout APCs. 0.5–1 million APCs are seeded in each 96-well plate. Engineered Jurkat T cells expressing a transmembrane-bound anti-IL-2 antibody to allow cytokine capture are co-transduced with exogenous TCRα/TCRβ pairs. (**D**) Flow cytometric analysis of T cells showing APC anti-CD69 versus PE anti-IL-2 (capture) after co-culture of TCR #1-expressing T cells with APCs expressing HLA-A*02:01 without or with an EBV CLGGLLTMV epitope-encoding gene. Data are representative of three independent experiments. (**E**–**F**) Schematic representation of T cell proximity labeling assay, which uses sequential staining to biotinylate APCs that present T cell-activating epitopes. Peptide-encoding genes are stably introduced into APCs expressing specific HLA genes. The APCs are co-cultured with T cells. T cells expressing TCRs that recognize the HLA–epitope complex express activation-induced markers on their cell surface. A biotin-labeled antibody stains the activation-induced markers. A streptavidin–HRP (horseradish peroxidase) fusion is linked to the biotinylated antibodies, to coat the activated T cell surface with enzyme. Biotin-xx-tyramide is added, and HRP causes promiscuous covalent biotin ligation in a proximity-dependent manner. Biotinylated APCs are sorted, and integrated epitope-encoding genes are amplified and sequenced (E). AIM markers or captured cytokines can be stained to label activated T cells (F). (**G**) Brightfield (left) and fluorescent (right) microscopy after APC/T cell co-culture. Proximity labeling was performed with an anti-CD2 antibody, and biotinylated cells were visualized using PE–streptavidin. T cells and interacting APCs became PE^+^, while APCs that were not in contact with T cells (marked in red asterisks) stayed PE^−^. (**H**) Brightfield (left) and fluorescent (right) microscopy of HLA-B*07:02- and CEF (32 CMV, EBV, or Flu epitopes) library-expressing APC clones (transduced at m.o.i. < 1) that were clonally expanded and then co-cultured with T cells expressing CMV-TCR #1 which targets the epitope TPRVTGGGAM. After co-culture, proximity labeling was performed with an anti-CD69 antibody, leading to staining of only specific APC/T cells. (**I**) Stained cells from (H) were sorted by MACS, and the epitope-encoding DNA was amplified and sequenced. Percent of total NGS reads in the sorted and unsorted populations are graphed, showing enrichment of the target epitope in the sorted population. (**J**) Difference between sorted cells and unlabeled cells in percent total reads for each epitope in (I). (**K**) Frequency of *z* score measurements for each epitope; *z* scores were calculated from the data in (J). (**L**) HLA-A*01:01-expressing APCs were transduced with the CMV genome library at a m.o.i. ∼ 5–10, and co-cultured with T cells expressing CMV-TCR #2 which targets the epitope YSEHPTFTSQY. After co-culture, proximity labeling was performed with an anti-CD69 antibody. Difference between sorted cells and unlabeled cells in percent total reads for each epitope are graphed. (**M**) Frequency of *z* score measurements for each epitope; *z* scores were calculated from the data in (L).

To determine whether a given TCR elicits T cell reactivity, we modified TCR knockout (KO) Jurkat T cells (*20*) to stably express a transmembrane-bound cytokine capture antibody, creating dual cytokine capture and AIM reporter T cells (**Fig. 1C**). We reasoned that staining for cytokine capture would provide a specific signal of T cell activation, but that the secreted cytokine could potentially be captured by nearby cells; while AIM expression, in our hands, was a less specific signal (*5*), but is cell-intrinsic. To test these dual reporter cells, we co-cultured TCR #1 (a recurrent TCR that is known to target the EBV epitope CLGGLLTMV from the LMP2A protein (*21*)) TCR-expressing T cells with APCs expressing HLA-A*02:01 alone, or HLA-A*02:01 and the target epitope in a minigene format (*20*). T cells co-cultured without epitope showed weak CD69 staining and negligible IL-2 capture, while T cells co-cultured with APCs expressing the cognate epitope showed strong CD69 induction and IL-2 capture (**Fig. 1D** and **fig. S1B**). The majority of the IL-2 capture (∼96%) occurred on CD69^+^ T cells (**Fig. 1D**), suggesting that the activated T cells capture their own IL-2 preferentially, while only a small fraction of IL-2 may leak to neighboring T cells.

### Establishing a system for T cell epitope identification using AIM

We reasoned that if we stained the activated T cell surface with antibodies linked to horseradish peroxidase (HRP) instead of PE (as in **Fig. 1D**), we could use HRP’s ability to catalyze proximity labeling (PL) (*22*) to send biotin molecules from the surface of the activated T cells down to the immunogenic antigen-presenting cells (**Fig. 1E**). This would allow us to sort the immunogenic peptides from our input libraries, and to pinpoint the peptides by next-generation sequencing (NGS) of the encoded peptides; and would also allow us to use the same endogenous signals of T cell activation – including AIM and cytokine secretion – that we used for screening, in order to preserve the sensitivity and specificity of these canonical readouts (*16, 23*).

After co-culture of T cells with APCs, we added biotinylated antibodies that bind to activation-induced markers (CD69) or to captured cytokines (IL-2). Second, we linked the biotin to a streptavidin–HRP (SA–HRP) fusion protein to coat the activated T cell surface with HRP enzyme. Third, we added the substrate biotin-xx-tyramide to enable HRP to covalently biotinylate membrane proteins at the T cell–APC interface (**Fig. 1, E** and **F**).

As a proof-of-principle of signal transfer from T cells to APCs, after co-culture, we stained T cells with a biotinylated anti-CD2 antibody (which labels all T cells), followed by staining with SA–HRP, biotin-xx-tyramide, and phycoerythrin–streptavidin (PE–SA). We confirmed by imaging that both the T cells and the interacting cells become biotinylated. APCs that were not in contact with T cells were not visibly biotinylated, consistent with localized signal transfer in a proximity-dependent manner (**Fig. 1G** and **fig. S1C**).

We then sought to determine whether our T cell proximity labeling method could separate T cell-activating APCs from non-activating cells in a pooled format (i.e. within a dish). As a proof-of-concept, we transduced a library of 32 CEF (CMV, EBV, and Flu) peptide-encoding oligonucleotides (oligos) (*20, 24*) into *HLA-B*07:02*–expressing APCs at an m.o.i. ∼ 1 and co-cultured the library with CMV-TCR #1 (a public CMV-reactive TCR with known specificity (*1, 25*)) TCR-expressing T cells. After performing T cell proximity labeling using an anti-CD69 antibody (**Fig. 1H** and **fig. S1D**), we sorted the PE^+^ APCs by magnetic activated cell sorting (MACS), isolated genomic DNA from the PE^+^ and PE^−^ cell populations, PCR amplified the peptide-encoding genes, and sequenced the amplicons using NGS. We found that 13.79% of total peptide-encoding reads encoded the CMV-TCR #1 target, TPRVTGGGAM in the positively-sorted (+sorted) cell population (**Fig. 1I**). In comparison, TPRVTGGGAM was represented by 1.44% of peptide-encoding reads in the negatively-sorted (–sorted) population (**Fig. 1I**). This difference of 12.35% was the most significant outlier amongst the library, with a *z* score of 5.23 (**Fig. 1, J** and **K**, and **table S5A**).

To determine whether we could identify a target epitope using a more complex library, we transduced our CMV library (4,867 encoded peptides) into *HLA-A*01:01*–expressing APCs and co-cultured the APC library with CMV-TCR #2 (another public CMV-reactive TCR with known specificity (*1, 22*)) TCR-expressing T cells. We found that 3.44% of total peptide-encoding reads contained the CMV-TCR #2 target – a YSEHPTFTSQY-containing 50-mer peptide – in the +sorted cell population. In comparison, this target was represented by 0.04% of peptide-encoding reads in the –sorted population. This difference of 3.40% was the most significant outlier amongst the library, with a *z* score of 48.78 (**Fig. 1, L** and **M** and **table S5B**).

These results demonstrated that our assay can identify a T cell-targeted epitope starting from a mixed pool of tens to thousands of peptide-encoding oligos.

### Systematic testing of putative antigen libraries for T cell reactivity

To screen for antigen reactivity, we had previously synthesized 4,867 peptide-encoding oligos that tile across all annotated genes of CMV (*20*), which has the largest genome amongst pathogenic human viruses (*26*). Given the high population incidence of EBV infection in the healthy U.S. population (*27*), and the observation that CMV and EBV may represent the most frequent known specificities amongst recurrent TCRs (*3*), we designed and synthesized 3,517 peptide-encoding oligos that tile across all annotated genes from three reference EBV strains. We also synthesized 1,266 peptide-encoding oligos that tile across all annotated genes from eight reference influenza A or B viral strains. Each of these oligos encodes a 50 amino acid peptide that overlaps with the adjacent encoded peptide (**Fig. 1C**). We cloned these oligo libraries into lentiviral expression vectors. We then seeded approximately 0.5–1 million HLA KO APCs (*20*) dispersed into a 96-well plate (i.e. approximately 5,000–10,000 APCs per well), and transduced them with our CMV, EBV, or Flu library at a multiplicity of infection (m.o.i.) of 5–10, along with each restricting HLA allele (**table S3**).

As a discovery set, we cloned 10 additional recurrent TCRα/TCRβ pairs (**fig. S1A** and **table S3**) into lentiviral expression vectors, and transduced the lentivirus into our engineered T cells. We co-cultured each of our TCR-transduced T cells with the APC libraries, and measured CD69 induction and IL-2 capture by flow cytometry. Given the complexity of these APC libraries, we expected that only a subset of the APCs express the cognate epitope. Accordingly, when T cells expressing TCR #1 were co-cultured with APCs transduced with CMV, EBV, or Flu peptide libraries, 11.2% of the T cells became dual CD69^+^/IL-2^+^ in response to the EBV library. In contrast, negligible CD69^+^/IL-2^+^ T cells were observed following co-culture with APCs presenting the CMV or Flu libraries, or with HLA-A*02:01 alone (**Fig. 2A**), demonstrating the specificity of the assay.

**Fig. 2.**
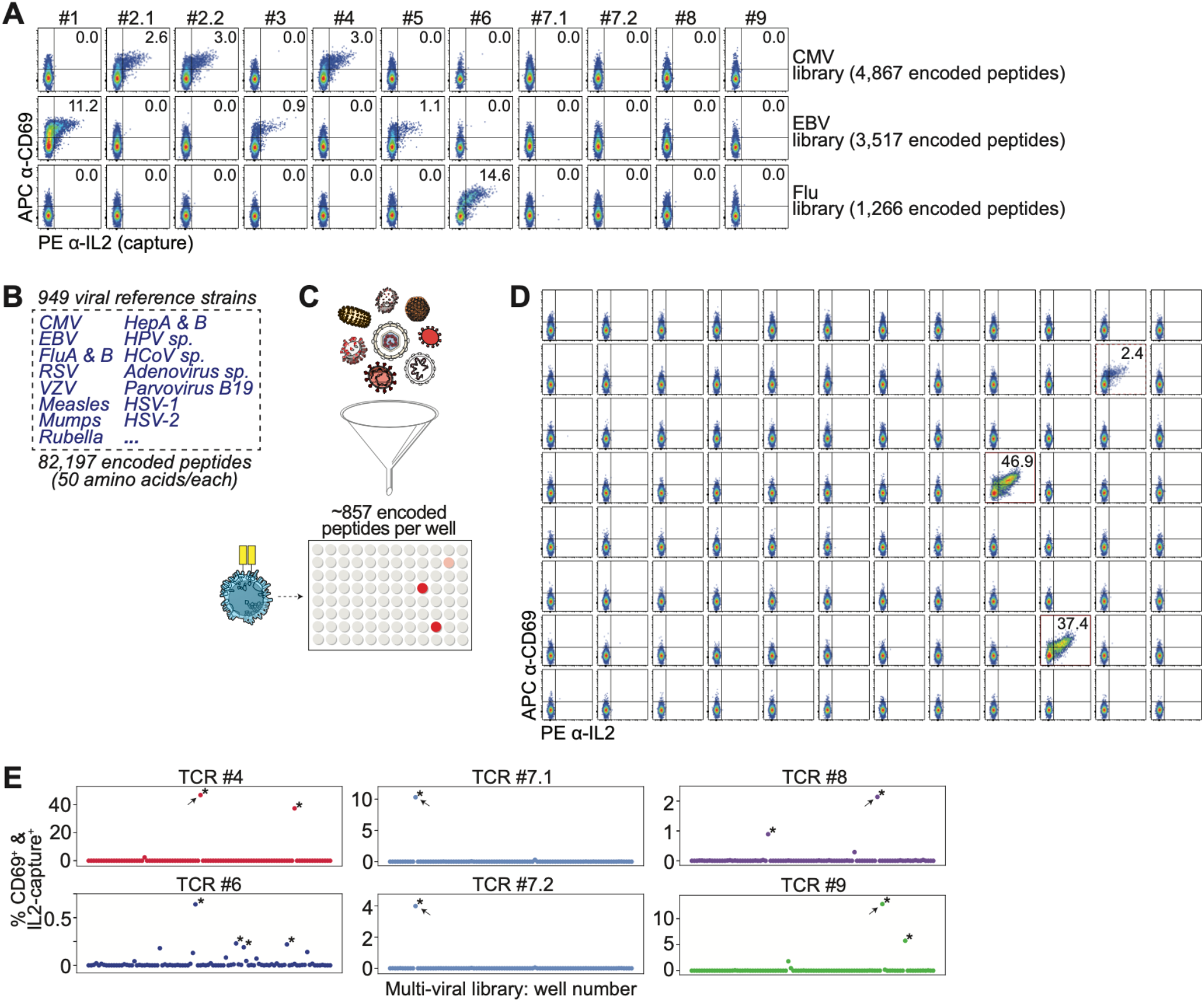
Screening libraries of reference human viruses. (**A**) Flow cytometric analysis of T cells showing APC anti-CD69 versus PE anti-IL-2 (capture) after co-culture of indicated TCR-expressing T cells with APCs expressing HLA-A*01:01, HLA-A*02:01, HLA-B*07:02, and HLA-B*08:01, with a genome-wide tiled CMV, EBV, or influenza encoded peptide library. (**B**) From 949 viral reference strains, 82,197 overlapping 50 amino acid peptide-encoding oligos were synthesized. (**C**) Schematic showing the 82,197 peptide-encoding genes partitioned into 96 sub-libraries of approximately 857 genes each, which were transduced into APCs and co-cultured with T cells. (**D**) Flow cytometric analysis of T cells showing APC anti-CD69 versus PE anti-IL-2 (capture) after co-culture of TCR #4-expressing T cells with APCs expressing HLA-A*01:01 and each sub-library in a 96-well plate. (**E**) Scatter plot of flow cytometry data plotting CD69^+^IL-2^+^ T cells in each well after co-culture of TCR-expressing T cells with APCs expressing each sub-library. Arrows point to wells that were used for T cell proximity labeling assays. \**z* score > 2.

TCR #2.1 and TCR #2.2 share the same TCRβ sequence but are paired with distinct TCRα chains (**table S2**). This TCRβ sequence was previously identified through pMHC tetramer sorting using HLA-A*01:01 loaded with the CMV-derived epitope VTEHDTLLY from the UL44 protein (25), although the corresponding TCRα chain was not determined in that study. We identified the TCRβ sequence in single-cell sequencing datasets, along with two possible paired TCRα heterodimers (*28, 29*). Consistent with the tetramer data, we confirmed that both TCRs elicit T cell activation in the presence of the CMV library but not the EBV or Flu libraries (**Fig. 2A**). We then validated that they target the CMV epitope VTEHDTLLY (**fig. S2A**). Similarly, the TCRβ #3 sequence was previously identified after pMHC tetramer sorting (*30*), but had an unknown TCRα heterodimer. After identification of the paired TCRα gene in a single-cell dataset (**table S2**), we confirmed that TCR #3 recognizes the EBV epitope RAKFKQLL from the BZLF1 protein (**fig. S2B**).

For the other TCRs, we could not find tetramer data supporting their target identity (*4*). After T/APC library co-culture, we found that 1 of these public TCRs (TCR #4) was reactive to our CMV library; 1 public TCR (TCR #5) was reactive to our EBV library; 1 public TCR (TCR #6) was reactive to our Flu library; and 4 public TCRs (TCRs #7.1, 7.2, 8, and 9) did not appear to be reactive to our CMV, EBV, or Flu libraries (**Fig. 2A**). For the CMV-, EBV-, and Flu-library reactive TCRs, we sought to test their reactivity to 32 common CMV, EBV, and Flu encoded peptides (i.e. the CEF32 peptide set (*20, 24*)). We determined that TCR #5 recognizes the EBV epitope RAKFKQLL from the CEF32 peptide set (**fig. S2, C** and **D**), but TCRs #4 and #6 did not appear to be reactive to the CEF32 set (**fig. S2C**).

Taken together, these results suggested that our workflow could efficiently determine if a TCR reacts to – or does not react to – a given viral library within a single well of a 96-well plate. Furthermore, we reasoned that if we could test thousands of encoded peptides in each well, we could theoretically screen a single TCR against hundreds of thousands of encoded peptides in a 96-well plate or in a subset of one plate.

### Screening a library of reference human viruses

To identify the target antigens of TCRs #7.1, 7.2, 8, and 9, we expanded our search to additional viral genomes. We thus synthesized 82,197 peptide-encoding oligos that tile across all annotated genes of 949 reference strains of viruses that are known to infect humans (*31*) (**Fig. 2B** and **table S4**); each of these oligos encodes a 50 amino acid peptide that overlaps with the adjacent encoded peptide (**Fig. 1C**).

As peptide library complexity increases, we reasoned that an increasingly smaller number of activated T cells would be detected. As a test, we co-cultured TCR #2.1-expressing T cells with APCs expressing *HLA-A*01:01* and (i) the target epitope, (ii) the 4,867 encoded peptide CMV library, or (iii) the 82,197 encoded peptide multi-viral library. As expected, the CD69^+^/IL2^+^ signal decreased as library complexity increased (**fig. S2E**). To improve sensitivity as well as to reduce the complexity of the epitope-containing library, we subdivided the full 82,197-peptide library into 96 smaller sub-libraries, each comprising approximately 857 peptide-encoding oligos. We transduced the sub-libraries into HLA-compatible APCs in a 96-well plate, and then co-cultured each T cell clone with each of the 96 sub-library wells (**Fig. 2C** and **fig. S2F**).

As a test, we used the CMV-reactive TCR #4-expressing T cell clone and the flu-reactive TCR #6-expressing T cell clone. For TCR #4, we identified 2 wells (with *z* score > 2) that contained activated T cells (**Fig. 2, D** and **E**, and **fig. S2C**). For TCR #6, we identified 4 wells (with *z* score > 2) that contained activated T cells (**Fig. 2E**). This was in line with our expectation: since the peptides are overlapping, the minimal HLA-binding peptide may be found within multiple 50-mer peptides; in addition, similar viral strains may contain the same minimal peptide but with slight differences in other regions of the 50-mer.

Extending this screening approach to our TCRs that were not reactive to CMV, EBV, or Flu (i.e. TCRs #7.1, 7.2, 8, and 9), we found that for each TCR, there were >/= 1 sub-library that activated the T cells (**Fig. 2E**). TCRs #7.1 and #7.2, which share the same TCRβ sequence, but are paired with different TCRα heterodimers, gave a similar pattern of sub-pool reactivity (**Fig. 2E** and **fig. S2G**), suggesting that they recognize the same epitope, but have allowance for sequence variability of the TCRα heterodimer (*25*).

These results suggested that we could narrow antigen specificity within a highly complex multi-viral library down to the ∼1% that contains the target epitope, and that TCRs #7.1, 7.2, 8, and 9 are indeed anti-viral TCRs, but target epitopes outside of CMV, EBV, and Flu.

### Determination of the target virus of public TCRs by T cell proximity labeling

Using T cell PL, we sorted the immunogenic encoded peptides using either the single virus library (**Fig. 2A**), or the most reactive sub-library in the multi-viral screen (**Fig. 2E**). After co-culture, staining, and APC sorting, we aliquoted a fraction of the +sorted and –sorted APCs back into cell culture. After multiple passages, we added fresh T cells to these APC aliquots, and found that the +sorted APCs caused increased T cell activation compared to the –sorted APCs (**Fig. 3A**), consistent with enrichment of the target epitope within the +sorted APCs. We PCR amplified the encoded peptides within the +sorted and –sorted APCs, performed NGS, and determined the encoded peptide sequence(s) enriched in the +sorted APCs.

**Fig. 3.**
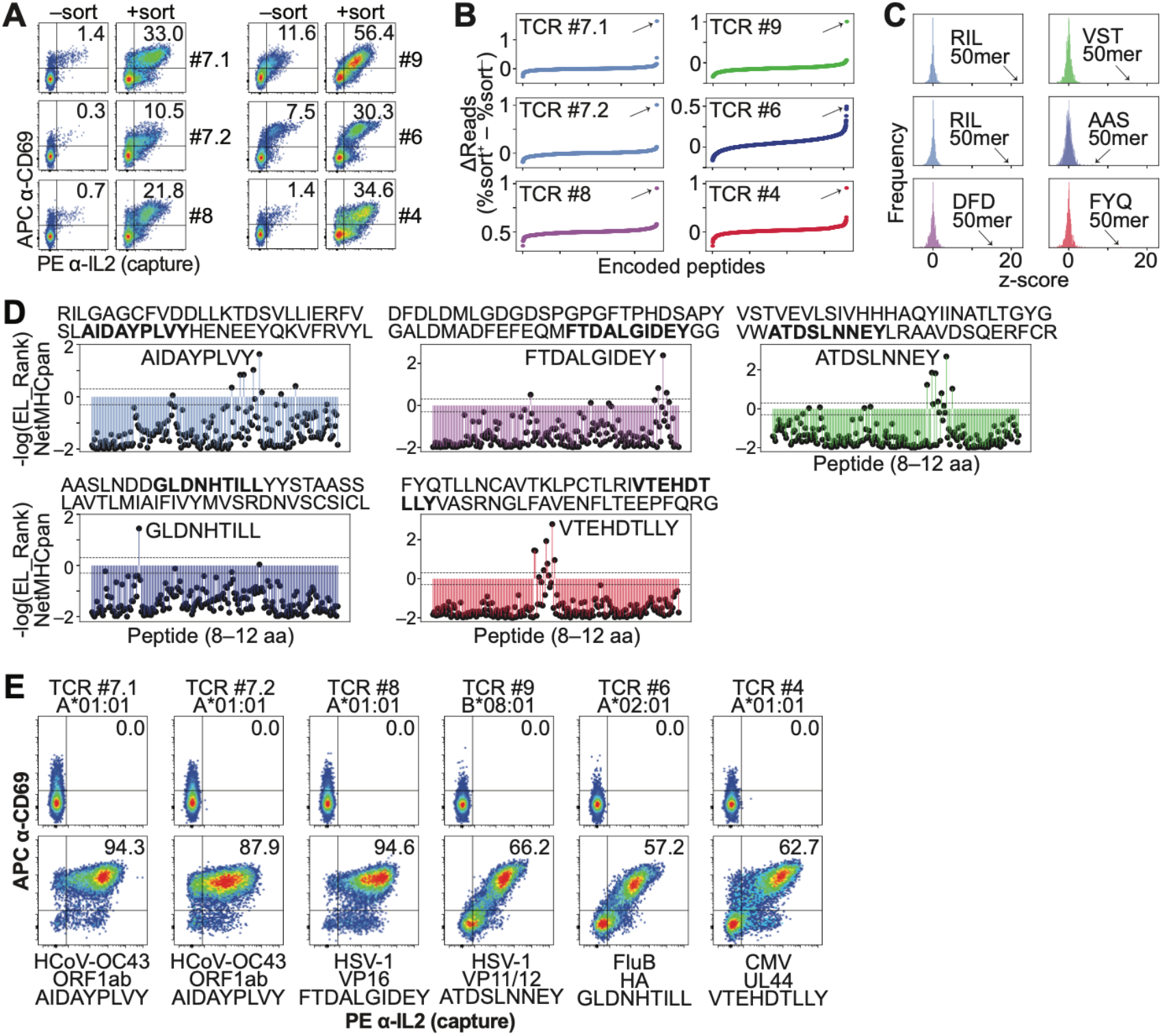
Determination of the target viral epitope using T cell proximity labeling. (**A**) Flow cytometric analysis showing APC anti-CD69 versus PE anti-IL-2 (capture) following T cell proximity labeling using the indicated TCRs and the target viral genome or the multi-viral sub-library. T cell clones were co-cultured with the unsorted or sorted APC populations, showing antigen enrichment within the sorted population. (**B**) Difference between sorted cells and unlabeled cells in percent total reads for each encoded peptide sequence after sequencing of the sorted and unsorted populations in (A). Arrows point to top candidate. (**C**) Frequency of *z* score measurements from screening data in (B). Arrows point to top candidate 50-mer encoded peptide. (**D**) %EL_Rank scores (−log) from NetMHCpan-4.1 for candidate 50-mer encoded peptide in (C). Thresholds for strong binders (SB) and weak binders (WB) are marked. (**E**) Flow cytometric analysis showing APC anti-CD69 versus PE anti-IL-2 (capture) after co-culture of indicated TCR-expressing T cells with APCs expressing the indicated restricting HLA allele without or with the candidate minimal epitope in (D). Data are representative of 1 to 3 independent experiments.

For both TCRs #7.1 and #7.2, using the most reactive sub-library (**Fig. 2E**), we identified a 50-mer (referred to here as the RIL 50-mer) from seasonal coronavirus OC43 as the top hit (**Fig. 3B, fig. S3A**, and **table S5, C** and **D**), with *z* scores of 21.97 and 19.79, respectively (**Fig. 3C**). The TCRβ #7.1 sequence is found in the peripheral blood of 14.4% (90/626) of individuals within the cohort (drawn from healthy donors in the U.S. (*1*)), and 46.2% of the HLA-A*01:01^+^ individuals (**fig. S1A** and **table S1**), consistent with the high (>90%) exposure rate of healthy individuals to seasonal coronavirus OC43, which causes the common cold (*6, 33*).

For TCR #8, using the most reactive sub-library (**Fig. 2E**), we identified a 50-mer (DFD) from HSV-1 (herpes simplex virus 1) as the top hit (**Fig. 3B** and **table S5E**), with a *z* score of 15.65 (**Fig. 3C**). The TCRβ #8 sequence is found within 5.0% (31/626) of the cohort, and 14.5% of the HLA-A*01:01^+^ individuals (*1*) (**fig. S1A** and **table S1**), consistent with the approximately 48% exposure rate of healthy individuals to HSV-1 (*9*). For the TCR #9, using the most reactive sub-library (**Fig. 2E**), we identified a 50-mer (VST) from HSV-1 as the top hit (**Fig. 3B** and **table S5F**), with a *z* score of 16.08 (**Fig. 3C**). The TCRβ #9 sequence is found within 5.0% (31/626) of the cohort, and 15.6% of the HLA-A*01:01^+^ individuals (*1*) (**fig. S1A** and **table S1**).

For TCR #6, using the Flu library (**Fig. 2A**), we identified the AAS 50-mer within influenza B as the top hit (**Fig. 3B** and **table S5G**), with a *z* score of 6.48 (**Fig. 3C**). The TCRβ #6 sequence is found within 5.1% (32/626) of the cohort, and 10.9% of the HLA-A*02:01^+^ individuals (*1*) (**fig. S1A** and **table S1**). For TCR #4, using the most reactive sub-library (**Fig. 2E**), we identified a 50-mer (FYQ) from CMV as the top hit (**Fig. 3B** and **table S5H**), with a *z* score of 13.47 (**Fig. 3C**). The TCRβ #4 sequence is found within 5.3% (33/626) of the cohort, and 10.2% of the HLA-A*01:01^+^ individuals (*1*) (**fig. S1A** and **table S1**).

Taken together, we identified viral targets for each of the public TCRs, including those with previously unknown target identity despite their common frequencies within healthy individuals. Our results revealed common viral targets apart from CMV, EBV, and flu A, thus providing a workflow to broaden existing knowledge of public TCR target specificities.

### Determination of complete TCR–peptide–HLA complexes

We then sought to molecularly define the complete TCR–peptide–HLA complexes containing each of the public TCRs. Because the libraries were screened using encoded peptides of 50 amino acids each, we predicted the minimal HLA-binding peptides using NetMHCpan-4.1 (*35*), and then functionally validated these peptide sequences.

For the coronavirus OC43-reactive TCRs #7.1 and #7.2, we functionally validated the RNA-dependent RNA polymerase (NSP12)-derived peptide AIDAYPLVY, which was the top ranked peptide within the RIL 50-mer predicted to bind to HLA-A*01:01 (**Fig. 3, D** and **E**, and **fig. S3A**). For the HSV-1-reactive TCR #8, we validated the alpha integrating protein (VP16)-derived peptide FTDALGIDEY, which was the top ranked peptide within the DFD 50-mer predicted to bind to HLA-A*01:01 (**Fig. 3, D** and **E**, and **fig. S3A**). For the HSV-1-reactive TCR #9, we validated the VP11/12-derived peptide ATDSLNNEY, which was the top ranked peptide within the VST 50-mer predicted to bind to HLA-A*02:01 (**Fig. 3, D** and **E**, and **fig. S3A**). For the influenza-reactive TCR #6, we validated the influenza B hemagglutinin (HA)-derived peptide GLDNHTILL, which was the top ranked peptide within the AAS 50-mer predicted to bind to HLA-A*02:01 (**Fig. 3, D** and **E**, and **fig. S3A**). For the CMV-reactive TCR #4, we validated the UL44-derived peptide VTEHDTLLY, which was the top ranked peptide within the FYQ 50-mer predicted to bind to HLA-A*01:01 (**Fig. 3, D** and **E**, and **fig. S3A**).

Taken together, these results demonstrated that our workflow can identify complete TCR– peptide–HLA complexes starting from TCRs of unknown target identity within the healthy T cell repertoire.

### Determination of cross-reactivity between viruses and species

We then sought to determine the extent of cross-reactivity of our TCRs against closely-related viruses and strains.

First, we aligned the coronavirus OC43 epitope, AIDAYPLVY, with the corresponding peptides within other seasonal coronaviruses (HKU1, NL63, and 229E) as well as within pandemic coronaviruses (severe acute respiratory syndrome (SARS)-CoV, Middle East respiratory syndrome (MERS), and SARS-CoV-2). These peptides – AIDAYPLDH, AIDAYPLSK, and AIDAYPLTK – were all predicted by NetMHCpan-4.1 to be weak binders to HLA-A*01:01, in contrast to the OC43 peptide which was predicted to be a strong binder (**Fig. 4A**). Testing of these peptides showed that the NL63/229E (AIDAYPLSK) and SARS-CoV-2 (AIDAYPLTK) peptides were able to activate TCR #7.1 – though the activation was reduced compared to the OC43 epitope – while the HKU1 (isolate N1, N2) (AIDAYPLVH) peptide did not activate, or very weakly activated this TCR (**Fig. 4A** and **fig. S4A**).

**Fig. 4.**
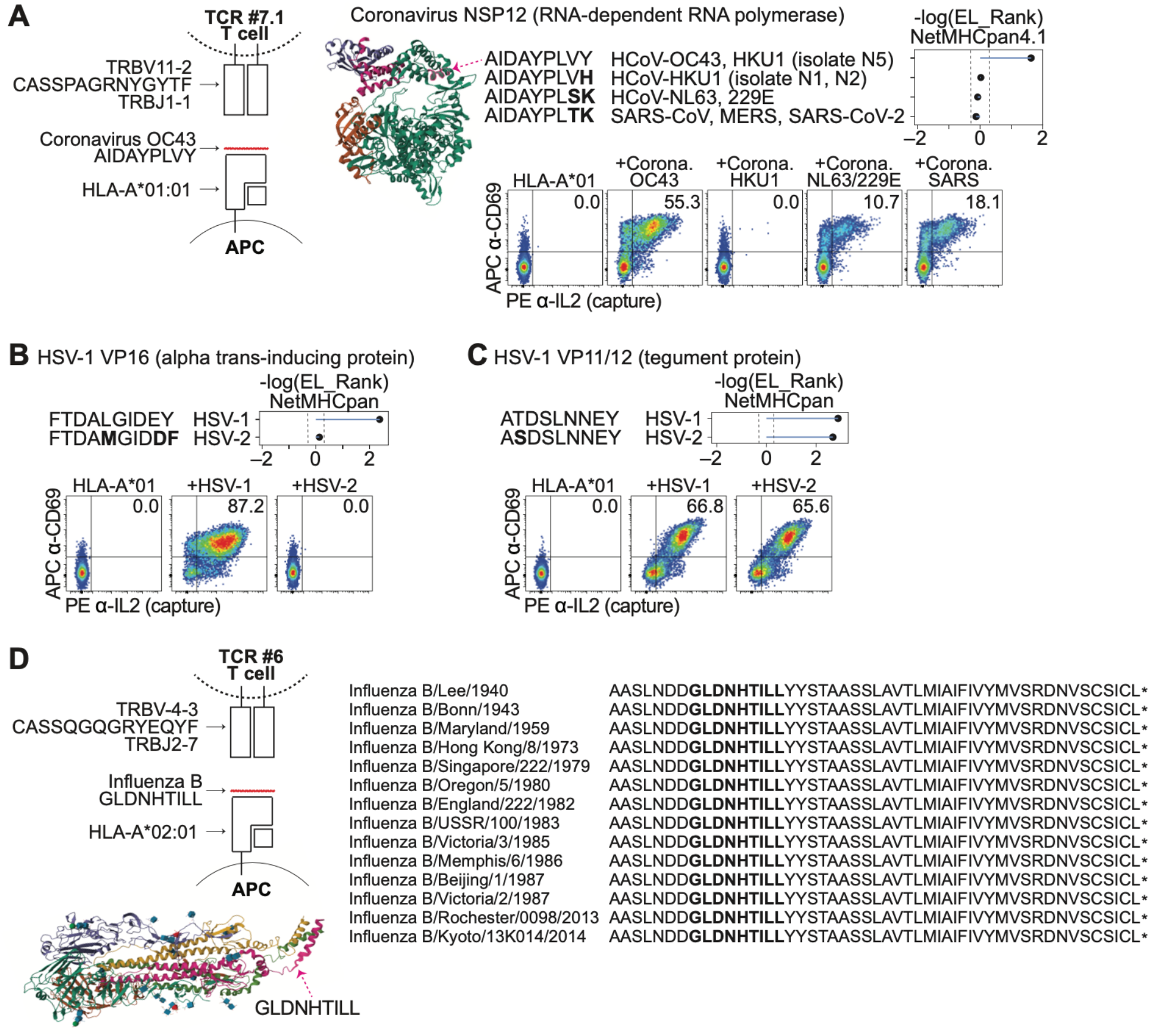
Determination of viral cross-reactivity. (**A**) Schematic representation of the complete TCR–peptide–HLA complex for TCR #7.1. Crystal structure of coronavirus NSP12 taken from PDB 6NUR; arrow points to the minimal epitope region. The coronavirus OC43 epitope as well as the corresponding peptides within HKU1, NL63/229E, and pandemic coronaviruses are shown, along with their %EL_Rank scores (−log) from NetMHCpan-4.1. Flow cytometric analysis showing APC anti-CD69 versus PE anti-IL-2 (capture) after co-culture of TCR #7.1-expressing T cells with APCs expressing HLA-A*01:01 without or with the panel of coronavirus peptides. Data are representative of three independent experiments. (**B**) The TCR #8 HSV-1 epitope as well as the corresponding peptide within HSV-2 are shown, along with their %EL_Rank scores (−log) from NetMHCpan-4.1. Flow cytometric analysis showing APC anti-CD69 versus PE anti-IL-2 (capture) after co-culture of TCR #8-expressing T cells with APCs expressing HLA-A*01:01 without or with the panel of coronavirus peptides. Data are representative of three independent experiments. (**C**) The TCR #9 HSV-1 epitope as well as the corresponding peptide within HSV-2 are shown, along with their %EL_Rank scores (−log) from NetMHCpan-4.1. Flow cytometric analysis showing APC anti-CD69 versus PE anti-IL-2 (capture) after co-culture of TCR #9-expressing T cells with APCs expressing HLA-A*01:01 without or with the panel of coronavirus peptides. Experiment was performed once. (**D**) Schematic representation of the complete TCR– peptide–HLA complex of TCR #6. Crystal structure of influenza hemagglutinin taken from PDB 6HJR; arrow points to the minimal epitope region. The minimal epitope sequences within the HA C-terminus from different influenza B strains are shown.

Similarly, we aligned the HSV-1 epitopes targeted by TCR #8 and TCR #9 with the corresponding peptides within HSV-2. For TCR #8, the HSV-1 peptide, FTDALGIDEY, was predicted by NetMHCpan-4.1 to be a strong binder to HLA-A*01:01, while the corresponding HSV-2 peptide, FTDAMGIDDF, was predicted to be a weak binder (**Fig. 4B**). For TCR #9, both the HSV-1 and HSV-2 peptides, ATDSLNNEY and ASDSLNNEY, were predicted to be strong binders to HLA-A*01:01 (**Fig. 4C**). Testing of FTDAMGIDDF suggested that TCR #8 (**fig. S4B**) did not cross-react with its counterpart in HSV-2 (**Fig. 4B**). In contrast, testing of ASDSLNNEY suggested that TCR #9 (**fig. S4B**) is cross-reactive with its counterpart in HSV-2 (**Fig. 4C**).

Finally, we aligned the influenza B epitope, GLDNHTILL, with the corresponding peptide within the strains of influenza B represented in our library. Notably, we did not identify any sequence variability between the strains (**Fig. 4D**). While the head and stalk regions of influenza hemagglutinin are known to be particularly variable between strains as they are common targets of neutralizing antibodies (*36*), the GLDNHTILL epitope is in the junction of the flexible linker region and the transmembrane domain (**Fig. 4D**), and may be less commonly subjected to antibody-mediated evolution (*37*).

Taken together, these results suggested that TCR #7.1 may be cross-reactive to other coronaviruses, including pandemic coronaviruses that emerged prospectively; TCR #8 appears specific to HSV-1, while TCR #9 appears to recognize both HSV-1 and HSV-2; and TCR appears to target all, or nearly all, influenza B strains.

## DISCUSSION

We describe a complete workflow to decipher public TCR reactivities *de novo*: (a) using our engineered dual reporter T cells, we demonstrate that TCRs can be rapidly tested against complete viral genomes in a single well of a 96-well plate, or against approximately 1,000 viral genomes (using 82,197 overlapping 50 amino acid encoded peptides) within one plate; (b) using T cell proximity labeling, we demonstrate that we can pinpoint each TCRs’ target epitope specificities. Using our AIMcap workflow, we identified specificities of previously “orphan” TCRs that are commonly circulating in healthy individuals. These TCRs are found in 3.5-14.4% of a U.S. cohort (**table S1**), suggesting that >10 million individuals in the U.S. have each of these T cell receptors in circulation. As hypothesized, we identified TCR specificities beyond common epitopes within CMV, EBV, and flu A (**fig. S4C**). For example, despite its high frequency as a T cell target, the seasonal (“common cold”) coronavirus epitope AIDAYPLVY was not previously found in existing epitope databases (*4, 36*); nor were there databased examples of many of our TCR– peptide–HLA complexes including our coronavirus OC43, HSV-1, and influenza B-reactive TCRs (*4*). Thus, we demonstrated that our strategy could successfully uncover public TCR specificities without starting with an *a priori* epitope panel.

Our AIMcap workflow enables efficient pre-screening of a panel of T cells for viral genome reactivity, prior to epitope identification. In combination with recently-developed high-throughput TCR cloning approaches (*37*), potentially thousands of TCRs could be tested in our workflow against the genomes of CMV and EBV, which may be the most common public TCR specificities (*3*). Furthermore, single viral genome libraries of other common public TCR specificities (e.g. other Herpesviridae or seasonal coronaviruses) could be synthesized and tested. To identify the underlying immunogenic APCs, we developed a novel T cell antigen identification technique based on proximity labeling using HRP (*22*) to label the APCs that present immunogenic peptide(s) to the T cell receptors. While other high-throughput antigen identification technologies have been described (*19*–*20, 38*), T cell proximity labeling is the first assay that initiates the signal from T cell cytokine capture or from an AIM signal, which are well-established endogenous signals of functional activation (*16, 23, 39*). Because cytokine secretion and AIM induction are not unique to CD8^+^ T cells, T cell proximity labeling should be readily modifiable (*20*) to study CD4^+^ T cell receptor specificities. Furthermore, the signal in T cell proximity labeling is generated using exogenous staining reagents that could be used without APC or T cell genetic engineering, and the HRP-linked antibody could readily be swapped out in order to bind to different targets (**Fig. 1F**).

Alternative methods to determine TCR target specificity include pMHC multimers (*15, 40*) and TCR clustering algorithms (*25, 41, 42*). While these are powerful tools, they require *a priori* knowledge of peptide-MHCs in their routine use. A combined approach which first utilizes our high-throughput workflow to identify immunodominant pMHCs in order to broaden pMHC databases beyond CMV, EBV, and Flu pMHCs (**fig. S4C**); followed by the development of pMHC multimers, their deployment to sort and sequence additional circulating T cell receptors, and the analysis of the sequencing results using TCR clustering may be the most efficient strategy to solve large numbers of public TCR specificities. Around 16,951 public HLA-associated TCRs have been enumerated, of which only a small fraction have known target identities (*3*). Gaining a more complete understanding of public TCR specificities has previously enabled the detection of past exposure to certain viruses with high sensitivity and specificity through deep TCR sequencing of peripheral blood samples (*1, 2, 12*). A more complete database of public TCR specificities may thus allow diagnosis of past exposure to a large number of antigens concurrently (*2*). Moreover, because certain viral exposures are linked to other acute/chronic diseases – including the association of influenza with Guillain-Barré syndrome (*43*) and the association of EBV with multiple sclerosis (*8, 9*) – identification of antigen-specific public TCRs may help shed light on these associations using patient cohort studies.

Intriguingly, we identified a public seasonal coronavirus-reactive TCR that appeared to be cross-reactive to pandemic coronaviruses that emerged prospectively after sample collection, suggesting that this TCR could have contributed to pre-existing immunity to pandemic coronaviruses (*33, 44–46*). However, while the OC43 peptide has strong NetMHC-predicted binding affinity, the corresponding SARS-CoV-2 peptide has weak predicted binding affinity to HLA-A*01:01 (**Fig. 4A**), suggesting that the physiological dose of the viral peptide within cells may influence its rate of recognition by T cells.

Along this same vein, despite the large potential space of HLA-binding peptides from each of these viruses (**fig. S3B**), several of the peptides that we identified through viral genome-wide screening have been repeatedly reported in databases (*4, 24, 36*). Our data are consistent with the hypothesis that many of these oft-studied epitopes are not just overrepresented in databases (*4*) due to ascertainment bias (*5*), but are at the top of viral immunodominance hierarchies (*47, 48*). Immunodominance is believed to be associated with strong HLA binding affinity, although a more complete model incorporates additional factors that make an epitope more “presented” (i.e. more likely to be seen by and activate a T cell receptor in an individual) than other peptides (e.g. the kinetics of viral protein expression, viral latency states, and viral load) (*47, 48*). In line with this model, we found that several of the epitopes that we identified – including the coronavirus and HSV-1 epitopes – are not just NetMHC-predicted strong HLA binders, but are amongst the highest ranked predicted strong binders within their respective virus (**fig. S3B**). By extension, due to immunodominance, the discovery of new public viral epitopes within other human viruses may saturate without the need to screen all public HLA-associated TCRs; i.e., the universe of highly public viral epitopes is likely a finite, discoverable set. Furthermore, the public nature of our influenza B-reactive TCR (TCR #6) may relate to the conservation of its target epitope between strains, i.e. there may be a higher event rate of exposure to conserved epitopes. By extension, other conserved influenza epitopes may be identifiable through public TCR target identification, which may aid in vaccine design (*35*).

Taken together, we demonstrate that T cell reactivity screening followed by T cell proximity labeling is an efficient way to reveal T cell memories *de novo*. Our overall workflow is agnostic to whether the TCR is public or private; is likely modifiable (*20*) to study CD4^+^ T cell specificities; and may not be limited to studying viral specificities, i.e. an analogous system (*20*) could be envisioned using bacterial peptides (*49*), autoimmune/alloimmune peptides, tumor peptides, and/or food/allergen peptides (*50*). Knowledge of TCR-peptide-HLA complexes is increasingly being used to develop diagnostics (*1, 2, 12*) and therapeutics (*51, 52*) such as vaccines and TCR-based cellular therapies (*13, 14*), underscoring the growing need for efficient strategies to solve HLA-peptide-TCR complexes *de novo*, as described herein.

## MATERIALS AND METHODS

### Cells, viruses, and reagents

Jurkat cells were obtained from ATCC, and maintained in RPMI (Gibco) supplemented with 10% FBS (Gemini Bio-Products) and penicillin–streptomycin (Gibco). HeLa and 293T cells were maintained in DMEM (Gibco) supplemented with 10% FBS. Self-inactivating minimal HIV-1 virus was produced in 293T cells using the vectors pLX301 or pLX303 (Broad Institute, Addgene plasmids #25895 and #25897), the packaging construct psPAX2, and the envelope plasmid pCMV-VSVG. Staining reagents were obtained from the following sources: PE-conjugated anti-IL-2 antibody (N7.48 A, Miltenyi), APC-conjugated and biotin-conjugated anti-CD69 antibody (FN50, BioLegend), biotin-conjugated anti-IL-2 antibody (B33-2, BD Biosciences), biotin-conjugated anti-CD2 antibody (RPA-2.10, BioLegend), streptavidin–HRP (BioLegend), biotin-xx-tyramide (SML3484, Sigma; or B40951, Thermo Fisher Scientific), PE–streptavidin (Thermo Fisher Scientific), and Zombie NIR viability dye (BioLegend).

### Recurrent TCR selection, cloning, and expression

TCRβ sequences that were clonally-expanded (defined as a frequency > 0.001) in peripheral blood of >1 healthy donor were identified in public TCRβ sequencing data (n = 626 donors with HLA class I typing) (*1*). TCRβ sequences that were recurrently found in donors typing for the same HLA allele were selected. A subset of these TCRβ sequences were re-identified in single-cell TCR sequencing datasets from solid tumors (*30, 31, 37, 53*–*56*), which allowed putative identification of the paired TCRα subunit. Paired TCRα and TCRβ sequences were cloned into pLX301 or pLX303. The TCR constant regions (*TRAC* and *TRBC1*) were modified with TRAC p.T48C and TRBC p.S57C mutations to facilitate TCR pairing (*22, 57*). TCR knockout Jurkat T cells (*20*) were transduced with *CD8A* and *CD8B* cDNAs; *LTBR* and *CARD11 p*.*D357N* cDNAs (*58, 59*); and a transmembrane-bound anti-IL2 antibody (*22, 60*). These engineered T cells were then co-transduced with lentivirus that contained the TCRα and TCRβ genes. Cloned TCR sequences are listed in table S3.

### Viral library construction

The CMV library was previously described (*20*). To construct the EBV library, we downloaded sequence data for all ORFs (open reading frames) that were annotated in the complete genome sequence of EBV strains AG876 and YCCEL1 in NCBI (accessions NC_009334 and AP015016). We added sequence data for all 92 ORFs that were annotated in the reference proteome of EBV strain B95-8 in UniProt (proteome ID UP000153037). To construct the influenza A/B library, we added sequence data for all ORFs that were annotated in the reference proteome of 6 influenza A strains (2 of H1N1, 3 of H3N2, and 1 of H7N7) and 2 influenza B strains in UniProt (proteome ID UP000009255, UP000171580, UP000096247, UP000166681, UP000115734, UP000123725, UP000099553, and UP000099653). To construct the multi-viral library, we downloaded all ORFs that were annotated in the UniProt database (*31*) with “Virus host” = “Homo sapiens (Human/Man) [9606]” and “Reviewed” = “Yes”. To construct the EBV and influenza tiled libraries, we tiled all encoded proteins with 50 amino acid peptides, each with a 32 amino acid overlap from the adjacent peptide, and added the C-terminal 50-mer. In total, this resulted in 3,517 and 1,266 unique peptides, respectively, after filtering out duplicates. To construct the multi-viral library, we tiled all encoded proteins with 50 amino acid peptides, each with a 25 amino acid overlap from the adjacent peptide, and added the C-terminal peptide. In total, this resulted in 82,197 unique peptides, after filtering out duplicates, from 949 viral strains. The peptide sequences were converted into DNA sequences, and PCR primer sequences were added to the 5’ and 3’ ends. The DNA libraries were synthesized as oligo pools (Twist Bioscience). Encoded peptides were cloned 3′ of a methionine and a spacer sequence (encoding HTVGLYM between the methionine and the encoded peptide to facilitate our cloning approach) into pLX301 using Gibson Assembly (New England BioLabs). For the multi-viral library, sub-libraries of about 857 DNA vectors/pool were propagated.

To express minimal peptide epitopes, peptide-encoding oligos (Yale Keck Oligo Synthesis Resource) were cloned in-frame 3’ of the human IL-2 signal sequence, and assembled into the lentiviral vector pLX301 using Gibson Assembly.

### Flow cytometry

HLA class I knockout (HLA KO) HeLa cells (*20*) were co-transduced with *HLA-A*01:01, HLA-A*02:01, HLA-B*07:02*, and *HLA-B*08:01* cDNAs. APCs were seeded in 96-well plates, and transduced with peptide-encoding DNA. The following day, T cells were added at a ratio between 8:1 and 16:1. After incubation at 37°C for 24 hours, cells were washed with DMEM. Cells were incubated with indicated antibodies for 30 minutes at room temperature, washed three times with phosphate-buffered saline (PBS; Gibco), and analyzed by flow cytometry (BD CytoFLEX).

### T cell proximity labeling assay

APCs and T cells were co-cultured as above. After incubation at 37°C for 24 hours, cells were gently washed with DMEM, and stained with a biotin anti-CD69 antibody, a biotin anti-IL2 antibody, or a biotin anti-CD2 antibody. Biotin-stained cells were gently washed with DMEM to remove excess antibody, and then stained with a streptavidin–HRP fusion. HRP-stained cells were gently washed with DMEM to remove excess enzyme, and then incubated with 1 mM hydrogen peroxide and 1 μM biotin-xx-tyramide for 10 minutes to enable proximity labeling. Biotin-stained cells were gently washed with DMEM. Cells were then dissociated with 0.25% trypsin-EDTA (Thermo Fisher Scientific), pooled, and stained with PE–streptavidin, APC anti-CD69 antibody, and Zombie NIR viability dye. Alternatively, the cells were stained with PE–streptavidin before dissociation, and imaged (Bio-Rad ZOE) to assess fluorescence. Live PE^+^ HeLa cells (+sort) were sorted from unlabeled cells (–sort) using FACS (Sony MA900) or alternatively, using a MACS Separator (Miltenyi) with anti-PE MicroBeads (Miltenyi). Aliquots of +sort and –sort HeLa cells were optionally placed back into a cell culture plate for expansion and repeat of co-culture to assess for enrichment of the target antigen.

### Amplification of encoded peptides and sequencing

Genomic DNA was extracted (DNeasy Blood & Tissue Kit; Qiagen) from PE-labeled cells (+sort) and from unlabeled cells (–sort). The integrated peptide-encoding DNA was amplified by PCR (Kapa HotStart ReadyMix; Roche) using barcoded primers complementary to sequences flanking the peptide-encoding sequences. NGS libraries were prepared from the PCR products and sequenced (Azenta Life Sciences).

### Data analysis

For each NGS read, the barcodes were identified to demultiplex the reads into associated samples, e.g. +sorted cells and –sorted cells. For each read, common sequences flanking the peptide-encoding sequence were identified, and the intervening peptide-encoding sequences were enumerated for each sample. For each sample, the fractional abundance of each encoded peptide was calculated. The difference in fractional abundance of each encoded peptide between pulldown and flow-through samples (annotated as % enrichment) was calculated and graphed.

Peptide–HLA binding predictions were performed using NetMHCpan-4.1 (*33*) using default parameters, including EL_Rank thresholds for strong binding (SB) peptides (0.5%) and weak binding (WB) peptides (2%).

### Statistical analysis

Data are reported as mean ± standard deviation. *P* values were calculated using Student’s t-test; \**P* < 0.05, \*\**P* < 0.005, \*\*\**P* < 0.0005. *Z* scores were calculated for encoded peptides identified in the +sort or –sort samples as the difference of each % enrichment value from the mean, divided by the standard deviation.

## Acknowledgments

We are grateful to members of the Lee laboratory for advice and valuable discussions. Funding: This work was supported in part by the National Cancer Institute of the National Institutes of Health (NIH) under Award Number NIH 5K08CA270191; by an Arthritis National Research Foundation (ANRF) grant; and by the Association for the Advancement of Blood & Biotherapies (AABB) Foundation Early-Career Scientific Research Grant. Competing interests: none. Data and materials availability: The published article includes all datasets generated in this study. Plasmids and cell lines generated in this work can be provided upon request to the senior author, subject to a material transfer agreement and barring any restrictions that apply to material owned by third parties.

