## Supplementary Figures for "*De novo* identification of the specificities of recurrent human T cell receptors"

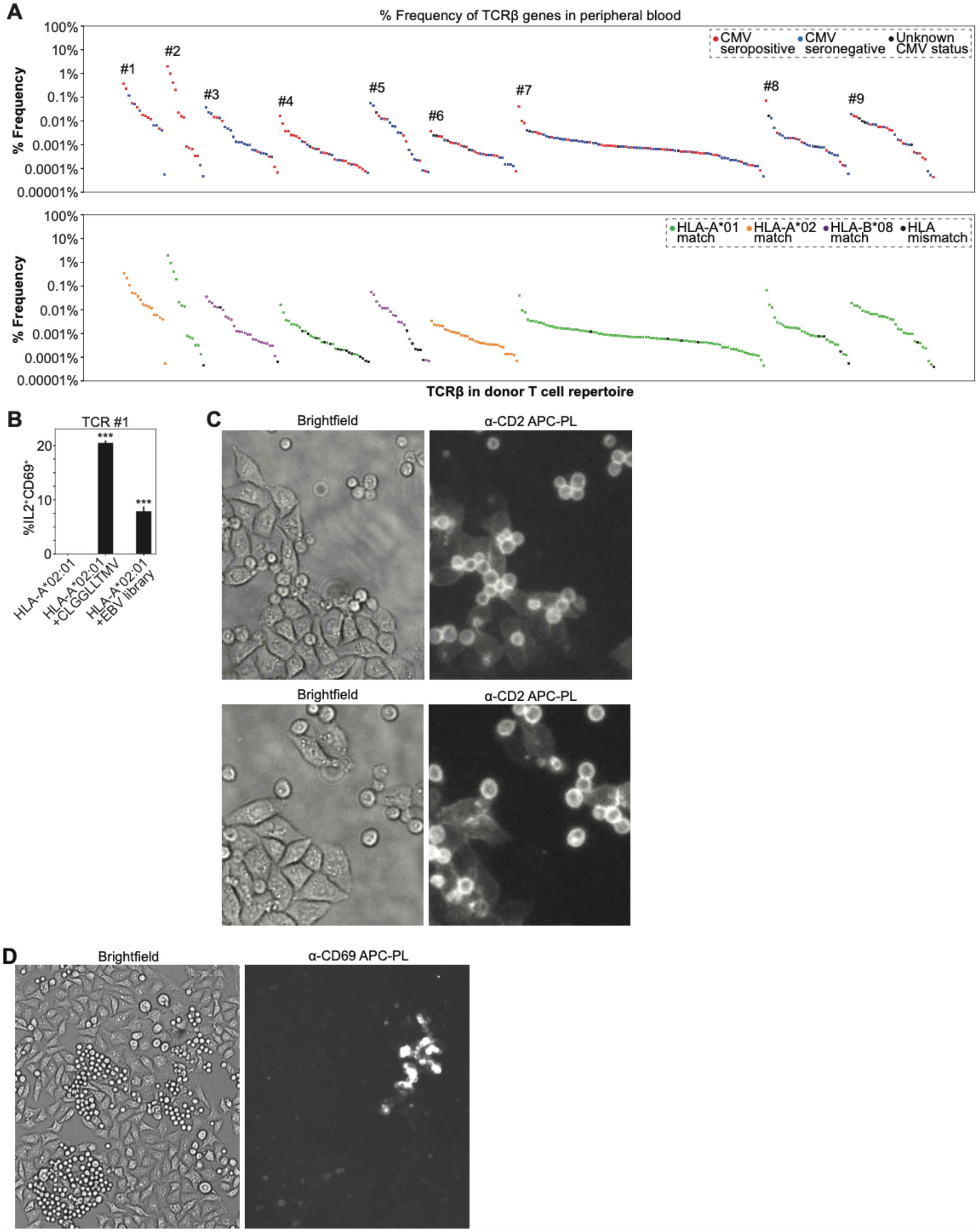


**Fig. S1. Development of the AIMcap system to identify the targets of public TCRs.** (**A**) Plot of % frequency (log scale) of specified TCRβ genes relative to all TCRβ genes sequenced in peripheral blood sample of donor. Each point represents an individual donor, and is colored according to CMV serological status (top) or according to HLA type (bottom). (**B**) Bar plot of

$CD69^+IL\text{-}2^+$ T cells after co-culture of TCR #1-expressing T cells with APCs expressing HLA-A*02:01 alone, with the CLGGLLTMV epitope, or with the exome-wide EBV library. ***$P <$ 0.0005 compared to cells expressing HLA alone (Student's t-test). Data are representative of two independent experiments. (**C**) Brightfield and fluorescent microscopy after APC/T cell co-culture. Proximity labeling was performed with an anti-CD2 antibody, and biotinylated cells were visualized using PE–streptavidin. T cells and interacting APCs became $PE^+$, while APCs that were not in contact with T cells stayed $PE^-$. (**D**) Brightfield (left) and fluorescent (right) microscopy of HLA-B*07:02- and CEF (32 CMV, EBV, or Flu epitopes) library-expressing APC clones (transduced at m.o.i. < 1) that were clonally expanded and then co-cultured with T cells expressing CMV-TCR #1 which targets the epitope TPRVTGGGAM. After co-culture, proximity labeling was performed with an anti-CD69 antibody, leading to staining of only specific APC/T cells.

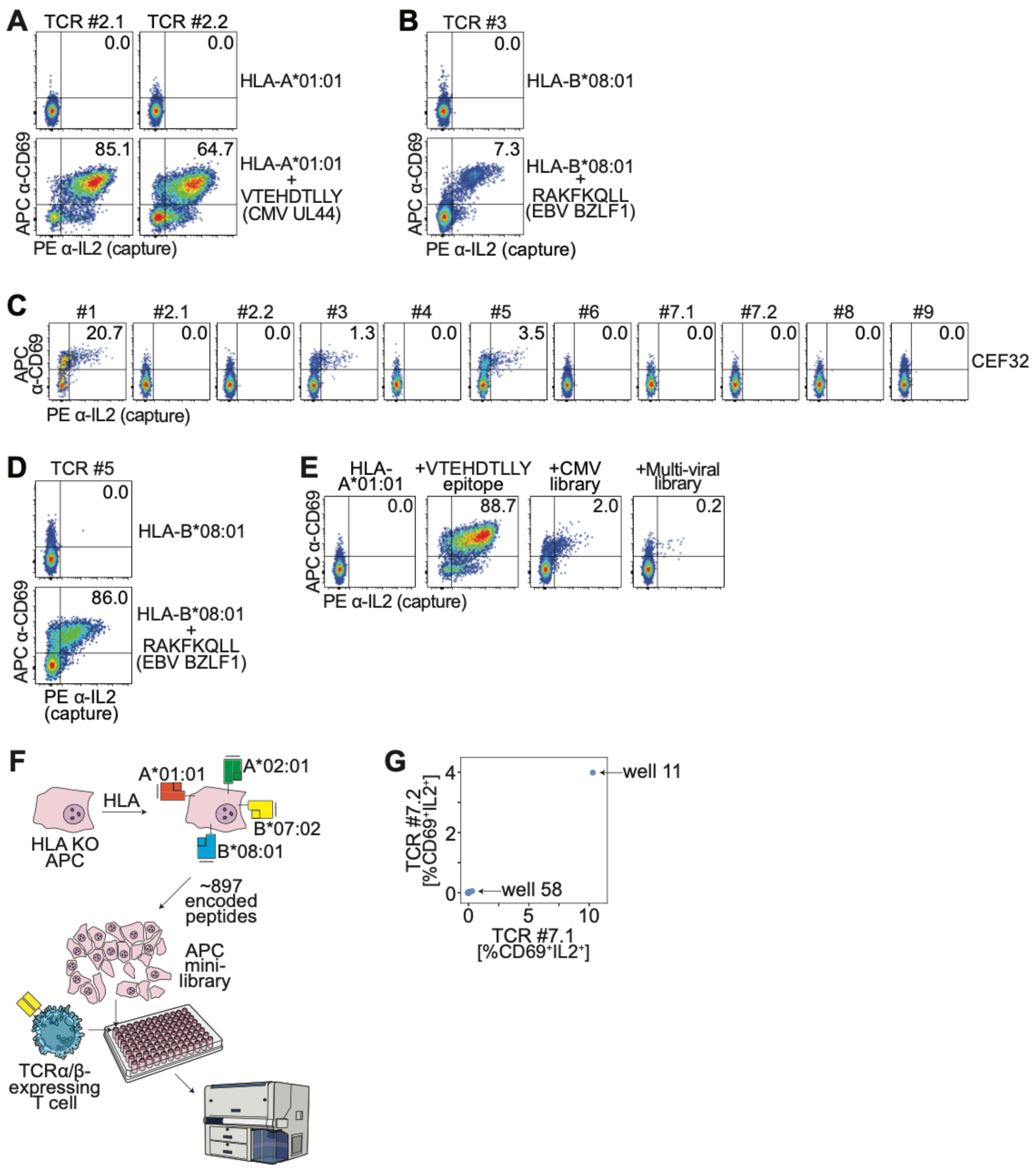


**Fig. S2. Screening libraries of reference human viruses.** (**A**) Flow cytometric analysis of T cells showing APC anti-CD69 versus PE anti-IL-2 (capture) after co-culture of T cells expressing TCR #2.1 or #2.2 with APCs expressing HLA-A*01:01 without or with the cognate CMV VTEHDTLLY epitope-encoding gene. Data are representative of three independent experiments. (**B**) Flow cytometric analysis of T cells showing APC anti-CD69 versus PE anti-IL-2 (capture) after co-culture of TCR #3-expressing T cells with APCs expressing HLA-B*08:01 without or with the cognate EBV RAKFKQLL epitope-encoding gene. Data are representative of three independent experiments. (**C**) Flow cytometric analysis of T cells showing APC anti-CD69 versus PE anti-IL-2 (capture) after co-culture of indicated TCR-expressing T cells with APCs expressing the restricting HLA allele with the CEF32 encoded peptide library. (**D**) Flow cytometric analysis of T cells showing APC anti-CD69 versus PE anti-IL-2 (capture) after co-culture of TCR #5-expressing T cells with APCs expressing HLA-B*08:01 without or with the cognate EBV RAKFKQLL epitope-encoding gene. Data are representative of three independent experiments.

(**E**) Flow cytometric analysis of T cells showing APC anti-CD69 versus PE anti-IL-2 (capture) after co-culture of TCR #2.1-expressing T cells with APCs expressing HLA-A*01:01 without or with the cognate CMV VTEHDTLLY epitope-encoding gene, the CMV exome library, or the multi-viral library. Data are representative of three independent experiments. (**F**) Schematic of the system to screen sub-libraries of the multi-viral library. HLA knockout APCs are transduced with HLA alleles. APCs in each well of the 96-well plate are transduced with a sub-library of the multi-viral library, and then co-cultured with TCR-expressing T cells. After co-culture, T cells are strained for activation-induced markers and flow cytometry is performed. (**G**) Comparison between TCRs #7.1 and #7.2 of %CD69$^{+}$IL-2$^{+}$ T cells in each well after co-culture of the indicated TCR-expressing T cells with APCs expressing each sub-library. Specific wells are labeled.

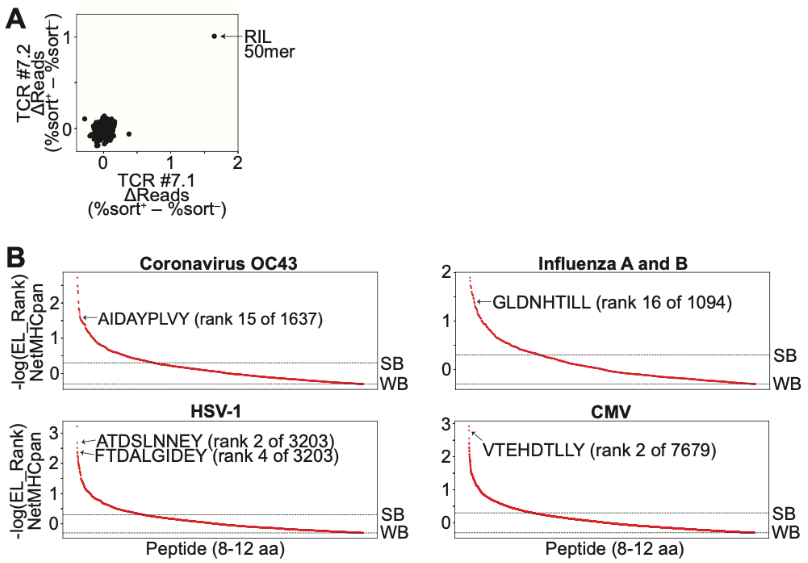


**Fig. S3. Determination of the target viral epitope using T cell proximity labeling.** (**A**) Comparison between TCRs #7.1 and #7.2 sequence after sequencing of the sorted and unsorted cells. (**B**) %EL_Rank scores (−log) from NetMHCpan-4.1 for all 8–12 amino acid peptides in indicated viruses from the multi-viral library. Thresholds for strong binders (SB) and weak binders (WB) are marked. Specific epitopes are labeled.

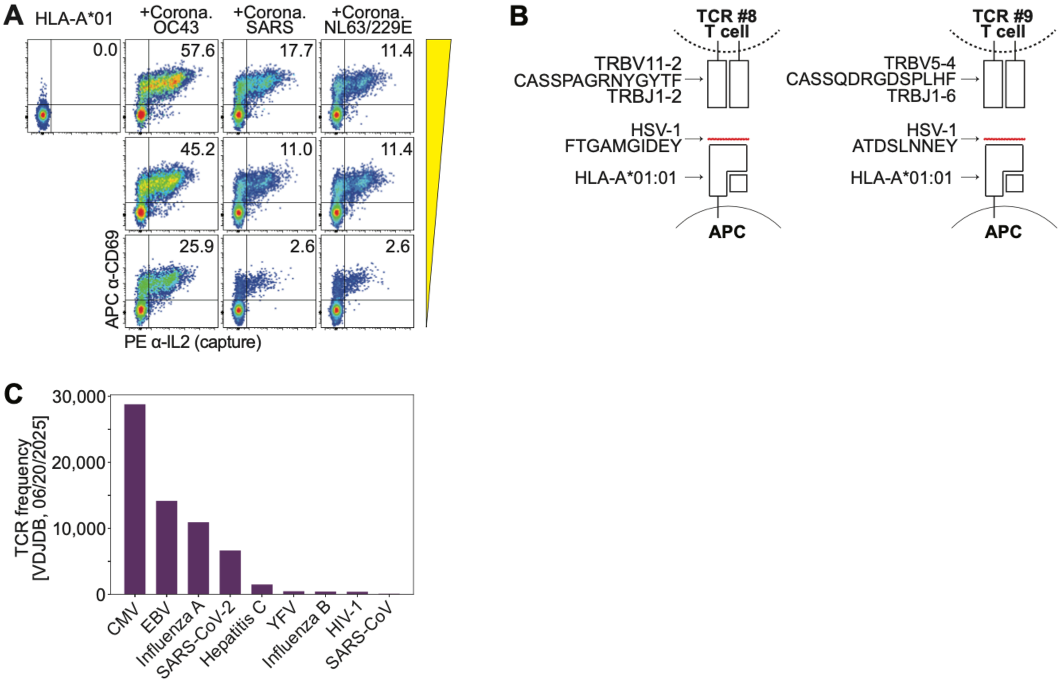


**Fig. S4. Determination of viral cross-reactivity.** (**A**) Flow cytometric analysis showing APC anti-CD69 versus PE anti-IL-2 (capture) after co-culture of TCR #7.1-expressing T cells with APCs expressing HLA-A*01:01 without or with a panel of coronavirus peptides from Fig. 5. A dose curve of lentivirus that contains each encoded peptides was used. (**B**) Schematic representation of the complete TCR–peptide–HLA complexes for TCR #8 and #9. (**C**) Bar plot of the frequency of TCRβ sequences in VDJdb (taken from 06/20/2025) that are denoted as reactive to the indicated viruses; the top 9 most frequently denoted viruses in VDJDB are shown, which represent 99% of the viral-reactive TCRβ sequences.
